# Selective deletion of interleukin-1 alpha in microglia does not modify acute outcome but regulates neurorepair processes after experimental ischemic stroke

**DOI:** 10.1101/2024.02.16.580635

**Authors:** Eloïse Lemarchand, Alba Grayston, Raymond Wong, Miyako Rogers, Blake Ouvrier, Benjamin Llewellyn, Freddie Webb, Nikolett Lénárt, Adam Denes, David Brough, Stuart M Allan, Gregory J Bix, Emmanuel Pinteaux

**Affiliations:** Division of Neuroscience, School of Biological Sciences, Faculty of Biology, Medicine and Health (FBMH), The University of Manchester, Manchester, UK; Geoffrey Jefferson Brain Research Centre, University of Manchester, Northern Care Alliance NHS Foundation Trust, The Manchester Academic Health Science Centre, Manchester, UK; Department of Neurosurgery, Clinical Neuroscience Research Center, Tulane University School of Medicine, Orleans, LA, USA; “Momentum” Laboratory of Neuroimmunology, HUN-REN Institute of Experimental Medicine, Budapest, Hungary

**Keywords:** ischemic stroke, microglia, interleukin-1 alpha

## Abstract

Inflammation is a key contributor to stroke pathogenesis and exacerbates brain damage leading to poor outcome. Interleukin-1 (IL-1) is an important regulator of post-stroke inflammation, and blocking its actions is beneficial in pre-clinical stroke models and safe in the clinical setting. However, the distinct roles of the two major IL-1 receptor type 1 agonists, IL-1α and IL-1β, and the specific role of IL-1α in ischemic stroke remain largely unknown. Here we show that IL-1α and IL-1β have different spatio-temporal expression profiles in the brain after experimental stroke, with early microglial IL-1α expression (4 h) and delayed IL-1β expression in infiltrated neutrophils and a small microglial subset (24-72 h). We examined for the first time the specific role of microglial-derived IL-1α in experimental permanent and transient ischemic stroke through microglial-specific tamoxifen-inducible Cre-loxP-mediated recombination. Microglial IL-1α deletion did not influence acute brain damage, cerebral blood flow, IL-1β expression, neutrophil infiltration, microglial nor endothelial activation after ischemic stroke. However, microglial IL-1α knock out (KO) mice showed reduced peri-infarct vessel density and reactive astrogliosis at 14 days post-stroke, alongside long-term impaired functional recovery. Our study identifies for the first time a critical role for microglial IL-1α on neurorepair and functional recovery after stroke, highlighting the importance of targeting specific IL-1 mechanisms in brain injury to develop more effective therapies.

## Introduction

Inflammation is a major contributor to stroke pathogenesis, and the role of inflammation driven by the pro-inflammatory cytokine interleukin-1 (IL-1) during post-stroke injury has been the focus of intense research. Indeed, pre-clinical studies have demonstrated the deleterious actions of IL-1 after stroke, whilst blocking its actions with the IL-1 receptor antagonist (IL-1Ra) is beneficial in experimental stroke (1) and in clinical setting (2). The cellular targets and actions of IL-1 in stroke are unclear, although IL-1 receptors (primarily the IL-1 type 1 receptor, IL-1R1) are expressed mainly on brain endothelial cells, and we have recently demonstrated, using cell-specific conditional gene deletion of IL-1R1, that brain endothelial cells and neurons are key targets of IL-1 in stroke (3). IL-1α and IL-1β are the two main agonists of the 11 members in the IL-1 family, and they are expressed centrally and peripherally after stroke (4, 5). The precise role and mechanisms of action of IL-1α remain largely unknown However, early studies suggested that IL-1α and IL-1β may exert specific differential actions during inflammation (6, 7) and, more recently, these isoforms were found to show a differential expression after experimental stroke, with early IL-1α expression in microglia preceding that of IL-1β (8). Furthermore, our recently published work found that IL-1α, but not IL-1β, selectively triggers angiogenesis *in vitro* (9) and IL-1α given at a low dose triggers angiogenesis and improves functional recovery after stroke (10), suggesting a specific role for IL-1α distinct to that of IL-1β after stroke.

Here we show that IL-1α is expressed by microglia early after stroke, followed by a later IL-1β expression in infiltrated neutrophils and a small microglial subset. Using a genetic mouse model allowing cell-specific deletion of IL-1α, we examined the specific contribution of microglial-derived IL-1α in ischemic stroke. Selective microglial IL-1α deletion did not modify acute outcome led to impaired neurorepair processes and long-term functional recovery after experimental ischemic stroke. These data reveal a potential critical role for microglial IL-1α in stroke and highlight the need for a much greater understanding of the respective contributions of IL-1 family members to cerebral ischemia, to allow the most efficient treatments to be developed.

## Methods

### Animals

All animal procedures were carried out in accordance with the Animals (Scientific Procedures) Act (1986), under a Home Office UK project license, approved by the local Animal Welfare Ethical Review Board, and experiments were performed in accordance with ARRIVE (Animal Research: Reporting of In Vivo Experiments) guidelines (11), with researchers blinded to genotype.

Animals were housed at 21±1°C, 55±10% humidity on a 12h light-dark cycle in Sealsafe Plus Mouse individually ventilated cages (Techniplast, Italy). Mice were supplied with Sizzle Nest nesting material (Datesand Ltd., UK) and cardboard enrichment tubes (Datesand Ltd., UK), and had *ad libitum* access to standard rodent diet (SDS, UK) and water. Mice were acclimatized to the facility for at least 1 week before the commencement of experimental work.

Experiments were performed on a total of 21 C57BL/6NCrl male mice (Charles River, UK) and on a total of 104 C57BL/6J male mice from an in-house colony (IL-1α^fl/fl^ and IL-1α^fl/fl^:Cx3cr1-Cre^ERT2^) at the University of Manchester. Complete schemes of the experimental designs are presented in Figures 1 to 4 and Figures S1 to S3. A total of 18 mice were excluded: 11 due to failure/complications of the experimental procedures, 1 due to mortality after intervention and 6 culled due to declining post-stroke health issues. Experiments using the in-house colony were performed on littermate controls. Microglial specific IL-1α knockout mice were generated by crossing mice in which exon 4 of the *IL1A* gene is flanked with loxP sites (IL-1α^fl/fl^) (12) with mice expressing CX3CR1 promoter-driven Cre recombinase (Cx3Cr1-Cre^ERT2^, JAX stock #020940) (13) that is expressed in the mononuclear phagocyte lineage. To induce Cre^ERT2^ activity and subsequent Cre-loxP-mediated deletion of IL-1α in CX3CR1-expressing cells, 3 to 6 months old mice were given tamoxifen by intraperitoneal injection for 5 consecutive days (2mg/100µL in corn oil, 75 mg/kg, Sigma-Aldrich). Control mice were Cre negative IL-1α^fl/fl^ treated with tamoxifen. Mice were recovered for 4 weeks after tamoxifen treatment to allow the peripheral myeloid wild type (WT) population to replenish, with sustained IL-1α deletion in microglia (14).

**Figure 1.**
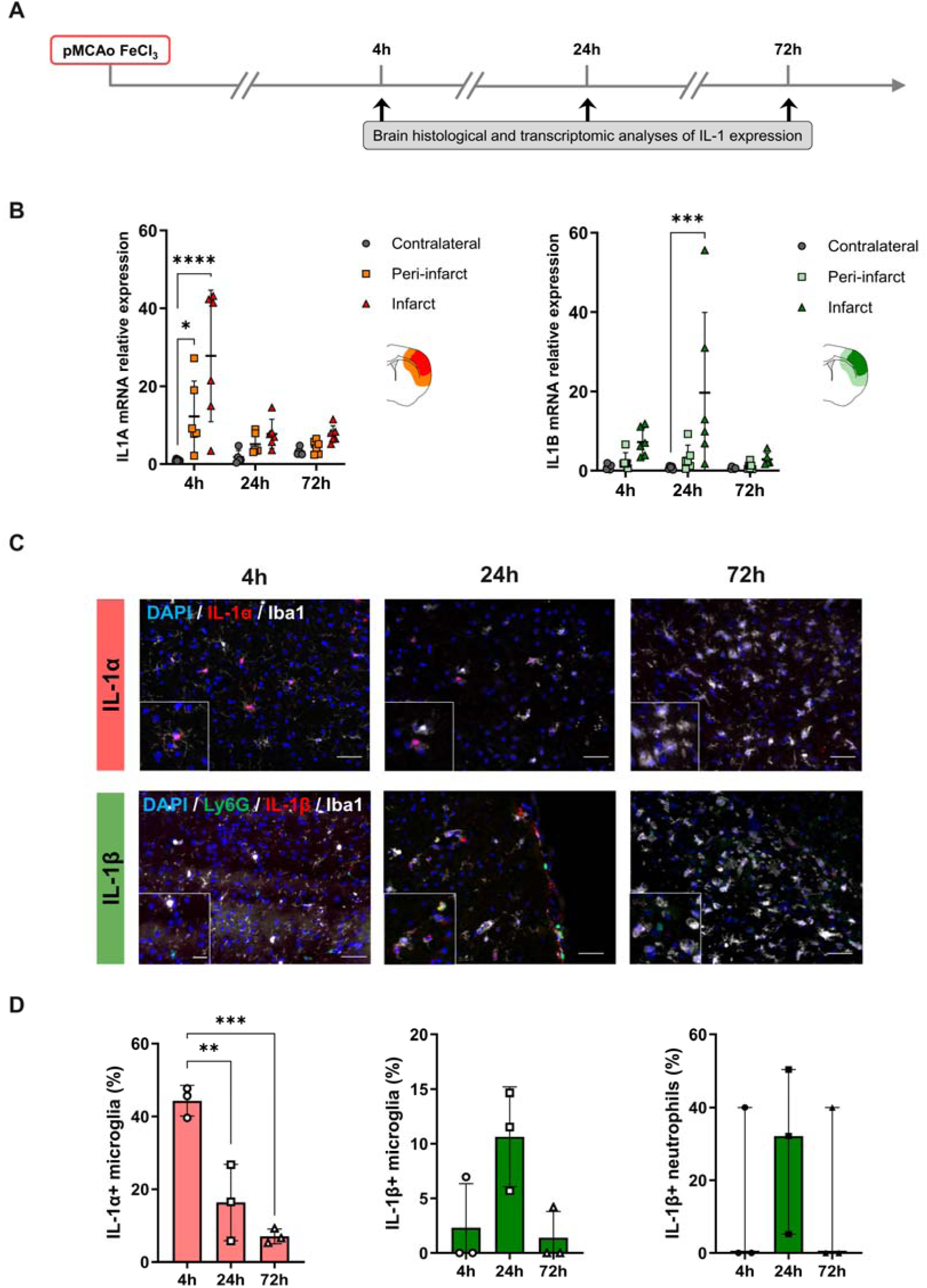
IL-1 brain expression during the early phase of distal pMCAo. (A) Schematic representation of the experimental design. (B) RT-qPCR analysis showing different spatio-temporal expression of *IL1A* (left) and IL1B (right) in the contralateral, peri-infarct and infarct areas at 4, 24 and 72 h after stroke (mRNA levels normalized to contralateral at 4 h); n=6/group, *p<0.05, ****p<0.0001 vs. contralateral, two-way ANOVA followed by Dunnet’s post hoc test. (C) Representative immunostaining of IL-1α (red), microglia (Iba1, white), neutrophils (Ly6G, green), IL-1β (red), and DAPI (blue) in the infarct at 4, 24 and 72 h after stroke. Scale bar in the large image is 50 µm and in the inset is 20 µm. (D) Percentage of IL-1α positive microglia in the infarct at 4, 24 and 72 h after stroke. (left; n=3/group, ***p<0.001, one-way ANOVA followed by Dunnet’s post hoc test). Percentage of IL-1β positive microglia and IL-1β positive neutrophils in the contralateral, peri-infarct and infarct areas at 4, 24 and 72 h after stroke. (right; n=3/group, one-way ANOVA or Kruskal-Wallis followed by Dunnet’s or Dunn’s post hoc test).

### Focal cerebral ischemia

Two different surgical models were used to induce a permanent or transient middle cerebral artery occlusion (pMCAo and tMCAo, respectively), both as clinically relevant models of focal ischemic stroke, the former recapitulating ischemic stroke with platelet-rich thrombi and unsuccessful reperfusion, and the latter mimicking large vessel occlusions followed by recanalization by endovascular thrombectomy.

Distal FeCl_3_-induced pMCAo was performed as a model of thrombotic cerebral ischemia, as described previously (15). In brief, mice were anesthetized with 4% isoflurane, placed in a stereotaxic device, and maintained under anesthesia with 2% isoflurane in a 70:30% mixture of N_2_O/O_2_. A small craniotomy (1 mm diameter) was performed on the parietal bone to expose the right MCA. A Whatman filter paper strip soaked in FeCl_3_ (40%, Sigma-Aldrich) was placed on the dura mater on top of the MCA for 2 times 5 min. Cerebral blood flow (CBF) in the MCA territory was measured continuously by laser Doppler flowmetry (Oxford Optronix, UK). A total of 11 mice were excluded from the study due to clot instability leading to inadequate occlusion and absence of ischemic lesion.

Transient focal cerebral ischemia was induced in two different research centers using the intraluminal model of proximal tMCAo, as previously described (3). Briefly, mice were anesthetized with isoflurane (4% for induction and 2% for maintenance in 30:70% mixture of N_2_O/O_2_, a midline incision was made on the ventral surface of the neck and the left common carotid artery isolated and ligated. A 6-0 monofilament (Doccol, Sharon, MA, USA) was introduced into the internal carotid artery via an incision in the common carotid artery (30 min MCAo) or in the external carotid artery (45 min MCAo). The filament was advanced approximately 10 mm into the common carotid with the filament making its way distal to the carotid bifurcation, to occlude the MCA. A 10_mm mark was made on the filament to visualize the required length to be inserted beyond the carotid bifurcation. There was an attrition of animals in the experimental groups followed up to 42 days after tMCAo, with 1 spontaneous death within the first 24 h, 4 animals culled on day 1, 1 culled at day 3 and 1 culled at day 4 relating to declining post-stroke health issues.

Body temperature was monitored and maintained at 37 ± 0.5 °C throughout both pMCAo and tMCAo procedures using a homoeothermic blanket and rectal probe (Harvard Apparatus, UK). Topical anesthetic (EMLA, 5% prilocaine and lidocaine, AstraZeneca, UK) was applied to skin incision sites prior to incision. Buprenorphine (0.05 mg/kg) was administered via subcutaneous injection at the time of surgery and at 24 h. Mice were weighed daily for the first week post-stroke, given mashed diet and administered with 0.5 mL saline daily until body weight stabilized (approximately at 4 days post-stroke).

### Laser Speckle Contrast Imaging (LSCI)

At 24 h after stroke induction, a laser speckle contrast imager (moorFLPI-2, Moor Instruments, UK) was used to measure CBF. Briefly, mice were anesthetized with isoflurane (4% for induction and 2% for maintenance in 30:70% mixture of N_2_O/O_2_) and fixed in a stereotaxic frame with ear bars and mouth bar to prevent movement. The scalp was exposed by a mid-line skin cut, and the skin was secured away using surgical clips to clear the skull at the region for imaging. An ultrasound gel was applied to the mouse skull and a glass coverslip was mounted on top to improve image quality. The CBF was then recorded by LSCI for 3 min (20 ms exposure time, 25 frame filter). Regions of interest (ROI) were drawn on the ipsilateral and contralateral brain hemispheres and analyzed using the moorFLPI Review V4 software (Moor Instruments) software. The percentage of CBF loss was calculated in the ipsilateral hemisphere compared to the contralateral brain hemisphere.

### Magnetic Resonance Imaging (MRI)

Mice underwent MRI scans at 4 h, 24_h and 8 d post-pMCAo. Animals were anesthetized with isoflurane (4% for induction and 2% for maintenance in 30:70% mixture of N_2_O/O_2_), and T2-weighted scans were conducted on Bruker Advance III console (Bruker Biospin Ltd, UK) using a 7 Tesla magnet. T2-weighted images were acquired using a multi-slice multi-echo (MSME) sequence: TE/TR 35 /7320 ms with 125 × 125 × 500 µm^3^ spatial resolution. Lesion volumes were measured using ImageJ software.

### Neurological deficit scoring

Mice were neurologically scored for focal deficits with the use of a 28-point neurological scoring system previously described (16). This is a cumulative score based on the following subcategories: body symmetry, gait, climbing, circling, front limb symmetry, compulsory circling, and whisker response. Animals are ranked from 0 (normal) to 4 (extreme deficit) for each subcategory. Neurological score was performed at 3, 7, 14, 21, 28 and 42 d post-stroke in the transient intraluminal MCAo model, which allows for a better assessment of behavioral deficits beyond the acute phase post-stroke, as opposed to distal cortical stroke models, which limit the investigation of long-term functional outcome.

### Tissue Processing

Anesthetized mice were transcardially perfused with cold saline followed by 50 mL of fixative (PBS 0.1 M, pH 7.4 containing 4% paraformaldehyde). Brains were removed, post-fixed for 24 h, cryoprotected (sucrose 30% in PBS; 24 h), and snap-frozen in isopentane at –30°C. Brains were then cut to a thickness of 30 μm using a freezing sledge microtome (Bright Instruments, UK) and then stored in cryoprotectant (0.05M Na_2_HPO_4_*2H_2_O, 5mM NaH_2_PO_4_, 30% anhydrous ethylene glycol, 20% glycerol) at –20°C until required, for free-floating immunohistochemistry, or embedded in optimal cutting temperature compound (OCT, PFM Medical UK, Ltd) prior to snap-freezing, and cut to a thickness of 10 μm using a cryostat (Leica CM1950, UK) for on-slide immunohistochemistry.

### Cresyl Violet stain

Cresyl violet staining was performed to measure lesion volume after tMCAo, as previously described (17). Briefly, 30 μm-thick brain sections were stained with 1% cresyl violet, and cover-slipped with DPX mounting medium (06522, Sigma-Aldrich). For each brain, infarct volumes were measured on defined coronal sections across the whole brain, spaced at approximately 360_µm apart (using image J). Each defined coronal section, with its brain co-ordinates and lesion was integrated to estimate total lesion volume for each brain and corrected for oedema.

### Immunohistochemistry

Brain sections were washed in Dulbecco’s phosphate buffeted saline (DPBS) with 0.1% Tween20, incubated in heat-mediated antigen retrieval in Tris-EDTA pH 8.6 solution for 20 min in a water bath set to 95°C, and incubated in blocking solution (1% BSA and 5% donkey serum in DPBS with 0.05% Tween20, 0.1% Triton X-100 and 0.2M glycine) for 1 h at room temperature. Sections were then incubated with rat–anti-Lymphocyte antigen 6 complex locus G6D (Ly6G) (1.25 μg/mL on-slide, 0.67 μg/mL free-floating, 127602; Biolegend), anti-Ionized calcium-binding adaptor molecule 1 (Iba1) (2 µg/mL on-slide, 1 µg/mL free-floating, ab178846, Abcam), goat–anti-IL-1α (1 μg/mL, AF-400-NA; R&D Systems), goat–anti-IL-1β (1 μg/mL, AF-401-NA; R&D Systems), goat–anti-Intercellular Adhesion Molecule 1 (ICAM-1) (1 μg/mL, AF796; R&D Systems), and chicken-anti-glial fibrillary acidic protein (GFAP) (1 μg/mL, Antibodies, ab4674) antibodies overnight at 4°C (in 1% BSA and 0.3% Triton X-100 in PBS). The following secondary antibodies were incubated for 2 h at room temperature: Alexa Fluor 488 donkey anti-rat, goat anti-chicken and donkey anti-goat (10 µg/mL; A21208, A11039, A32814), Alexa Fluor 568 donkey anti-rabbit (10 µg/mL; A10042), and Alexa Fluor 647 donkey anti-rat, donkey anti-rabbit (10 µg/mL; A48272, A31573). IL-1β and IL-1α signal was amplified with Tyramide SuperBoostTM (Thermo Fisher) using biotinylated horse–anti-goat IgG (7.5 µg/mL, BA-9500; Vector) secondary antibody. To assess vessel density, brain slices were stained with DyLight-488 conjugated tomato lectin (1:200, ThermoFisher, L32470) during secondary antibody incubation. Sections were then washed again and counterstained with DAPI (1:20 in diluted H_2_O) to stain for cell nuclei, after which they were then washed for a final time and mounted with Prolong Gold antifade reagent (Thermo Fisher Scientific; P36934).

Images were collected on Zeiss Axioimager.M2 upright microscope using a 20X Plan Apochromat objective and captured using a Coolsnap HQ2 camera (Photometrics) through Micromanager software (v1.4.23), or on a 3D-Histech Pannoramic-250 microscope slide-scanner using a x20/0.30 Plan Achromat objective (Zeiss, Germany) and captured using the Case Viewer software (3D-Histech, Hungary). Quantitative analysis was performed on 3-6 low magnification images of the lesion taken from at least 3 different coronal brain slices. Images were processed using ImageJ. Microglial activation based on morphology (Iba-1+ cells), neutrophil infiltration (Ly6G+ cells), vessel density (lectin-FITC+) and vessel activation (ICAM-1+) were analyzed using the Ilastik software program (18). To train the pixel classification algorithm, labelling was performed on at least 6 randomly selected images. All color/intensity, edge, and texture features were included, with a σ0 =□0.30, σ1 =□0.70, σ2 =□1.00, σ3 =□1.60, σ4 =□3.50, σ5 =□5.0 and σ6 = 10.0.

### Real-Time Quantitative Polymerase Chain Reaction

Total RNAs were extracted from samples with TRIzol Reagent (Thermo Fisher) according to the manufacturer. RNA (1 µg) was converted to cDNA using Super Script III Reverse Transcriptase (Thermo Fisher). Real-Time Quantitative polymerase chain reaction (RT-qPCR) was performed using Power SYBR Green PCR Master Mix (Thermo Fisher) in 384-well format using a 7900HT Fast Real-Time PCR System (Applied Biosystems). Three microliters of 1:20 diluted cDNA was loaded with 200 mmol/L of primers in triplicate. Data were normalized to the expression of the housekeeping gene Hmbs. Specific primers were designed using Primer3Plus software (http://www.bioinformatics.nl/cgi-bin/primer3plus/primer3plus.cgi). Primers used were: Il1b Forward— AACCTGCTGGTGTGTGACGTTC, Il1b Reverse—CAGCACGAGGCTTTTTTGTTGT, Il1a Forward— TCTCAGATTCACAACTGTTCGTG, Il1a Reverse—AGAAAATGAGGTCGGTCTCACTA. Expression levels of genes of interest were computed as follows: relative mRNA expression=E^−(Ct^ ^of^ ^gene^ ^of^ ^interest)^/ E^−(Ct^ ^of^ ^housekeeping^ ^gene)^, where Ct is the threshold cycle value and E is efficiency.

### Statistical analysis

Statistical analyses were performed using GraphPad Prism 8.0. All values are expressed as mean ± standard deviation (SD) or median (InterQuartile Range, IQR) according to the normal or non-normal distribution of the represented variable, respectively. The normality of continuous variables was assessed using the Shapiro-Wilk test (n < 30) or Kolmogorov-Smirnov test (n ≥ 30). Data were analyzed using unpaired t-test, one-way ANOVA (followed by Dunnet’s multiple comparisons post hoc test), two-way ANOVA (followed by Dunnet’s or Sidak’s multiple comparisons post hoc test), or by fitting a mixed effects model using Restricted Maximum Likelihood (REML), followed by Sidak’s multiple comparisons post hoc test. The Kruskal Wallis test (followed by Dunn’s multiple comparisons post hoc test) was used for non-normally distributed variables. The significant level was set at p<0.05.

## Results

### IL-1**α** and IL-1**β** are differentially expressed during the acute phase of ischemic stroke

Gene expression of *IL1A* and *IL1B* was analyzed by RT-qPCR in the infarct, peri-infarct and contralateral brain cortical areas dissected from mice subjected to distal pMCAo at 4, 24 or 72 h post-stroke (Fig. 1A). The expression of *IL1A* was affected both by time after pMCAo (p=0.0003) and the dissected brain regions (p=0.0001). A significant increase of *IL1A* mRNA was observed in the peri-infarct and infarct areas at 4 h after stroke (12-fold and 28-fold increase, p=0.0116 and p=0.0001 vs. contralateral, respectively); Fig. 1B. The expression of *IL1B* was also affected both by time after pMCAo (p=0.0339) and the dissected brain regions (p=0.0007). However, *IL1B* mRNA expression was increased later at 24 h post-stroke in the infarct areas (7-fold increase, p=0.0001 vs. contralateral); (Fig. 1B). To assess the protein expression profile of IL-1 after stroke and identify the cell types expressing IL-1α and IL-1β, we performed immunostaining for both cytokines on brain sections. IL-1α was detected exclusively in microglia (Iba1+, IL-1α+), with a significantly greater expression at 4 h compared to 24 and 72 h. Specifically, 44% of microglial cells were IL-1α+ at 4 h, compared to 16% and 7% at 24 and 72 h post-MCAo, respectively (p=0.0037 and p=0.0008 vs. 24 and 72 h, respectively); (Fig. 1C-D). However, IL-1β expression peaked at 24 h and was observed only in a small subset of microglia (11% at 24 h vs. 2% and 1% at 4 and 72 h post-MCAo respectively), although not reaching statistical significance between time points (p=0.0626 and p=0.9372 at 24 h vs. 4 and 72 h post-MCAo, respectively). In contrast, IL-1β was mostly expressed by neutrophils (Ly6G+), peaking at 24 h post-MCAo (29% vs. 13% at 4 and 72 h post-MCAo), again with no differences between timepoints (p=0.4815 and p=0.9999 at 24 h vs. 4 and 72 h post-MCAo, respectively. However, we did not observe significant changes between the different timepoints (p>0.05 for all comparisons; Fig. 1C-D).

### Successful microglial IL-1**α** deletion does not affect acute ischemic stroke outcome

Based on our previous results showing that microglia are the major source of brain IL-1α acutely after stroke, we targeted microglial-derived IL-1α to investigate its specific role after stroke. Successful deletion of IL-1α in microglia was confirmed by immunohistochemistry in IL-1α^fl/fl^:Cx3cr1-Cre^ERT2^ mice subjected to distal pMCAo at 4 h after stroke, which showed no IL-1α+ staining as opposed to control IL-1α^fl/fl^ mice, which did show IL-1α+Iba1+ staining (Fig. S1).

First, we investigated the influence of microglial IL-1α depletion on acute stroke outcome, at 24 h after both distal pMCAo and proximal tMCAo (Fig. 2). At 24 h after pMCAo, IL-1α^fl/fl^:Cx3cr1-Cre^ERT2^ mice showed no changes, neither in lesion volume (18.2±9.9 mm^3^ vs. 12.2±6.4mm^3^ in the IL-1α^fl/fl^ group, p=0.2060; Fig. 2A-B), nor in % of CBF loss in the ipsilateral vs. contralateral brain hemisphere (–19% vs. –14% in the IL-1α^fl/fl^ group, p=0.2991; Fig. 2A-B). In order to assess infarct evolution beyond 24 h after pMCAo, IL-1α^fl/fl^:Cx3cr1-Cre^ERT2^ mice and control IL-1α^fl/fl^ mice were followed for 8 days and underwent MRI to track infarct size longitudinally at 4 h, 24 h and 8 d post-MCAo by T2-weighted imaging (T2WI), again showing no changes in lesion volume associated to genotype at the different timepoints nor in lesion volume progression with time (p>0.05 for all comparisons vs. IL-1α^fl/fl^ group; Fig. 3). We also investigated whether deletion of microglial IL-1α could affect acute immune cell responses, at 24 h after pMCAo. As expected, an increase in microglial cell density was observed in the infarct areas in both IL-1α^fl/fl^:Cx3cr1-Cre^ERT2^ (p=0.0069 vs. contralateral, p<0.001 vs. peri-infarct), and control IL-1α^fl/fl^ mice (p=0.0069 vs. contralateral, p<0.001 vs. peri-infarct). An increase in neutrophil infiltration was also observed in the infarct areas in both IL-1α^fl/fl^:Cx3cr1-Cre^ERT2^ (p=0.0357 vs. contralateral, p=0.1099 vs. peri-infarct), and control IL-1α^fl/fl^ mice (p=0.0098 vs. Contralateral, p<0.001 vs. Peri-infarct). However, there were no changes between genotypes in neither microglia density nor neutrophil infiltration in the contralateral, peri-infarct and infarct areas (p>0.05 for all comparisons vs. IL-1α^fl/fl^); (Fig. S2). Moreover, IL-1β expression in both microglia and neutrophils remained unchanged between genotypes (p>0.05 for all comparisons vs. IL-1α^fl/fl^; Fig. S2).

**Figure 2.**
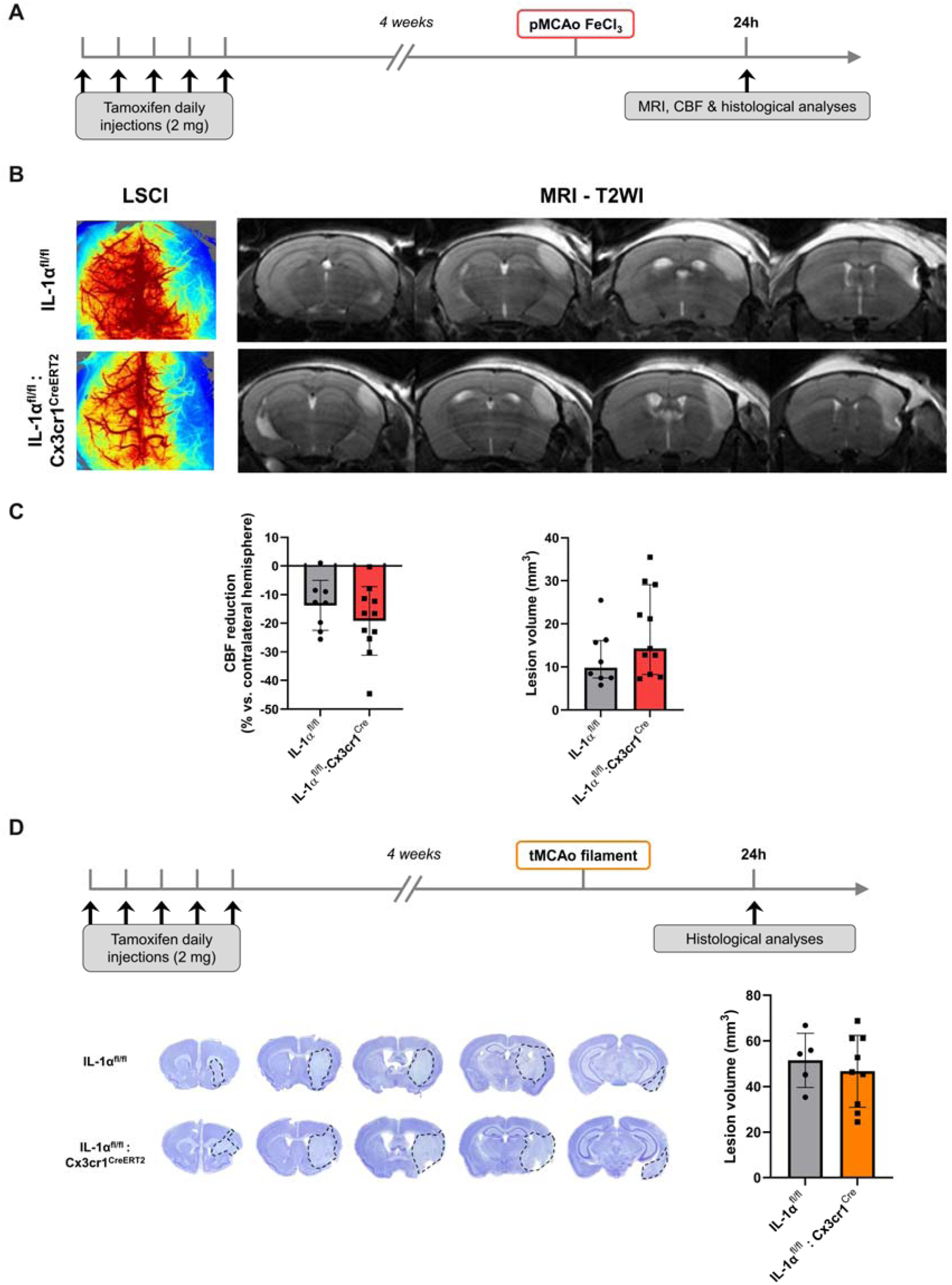
Microglial IL-1α deletion does not influence acute brain damage after pMCAo nor tMCAo. (A) Schematic representation of the experimental design to study acute outcome after permanent stroke. (B) Representative LSCI and CBF reduction quantification (left) and representative T2-WI and lesion volume quantification (right) at 24 h after pMCAo in IL-1α^fl/fl^ and IL-1α^fl/fl^:Cx3cr1-Cre^ERT2^ mice (n=8-11/group, unpaired t-test). (C) Schematic representation of the experimental design to study acute outcome after transient stroke. (D) Representative cresyl violet-stained brains (lesion delineated by dashed lines) and lesion volume quantification (n=5-9/group, unpaired t-test).

**Figure 3.**
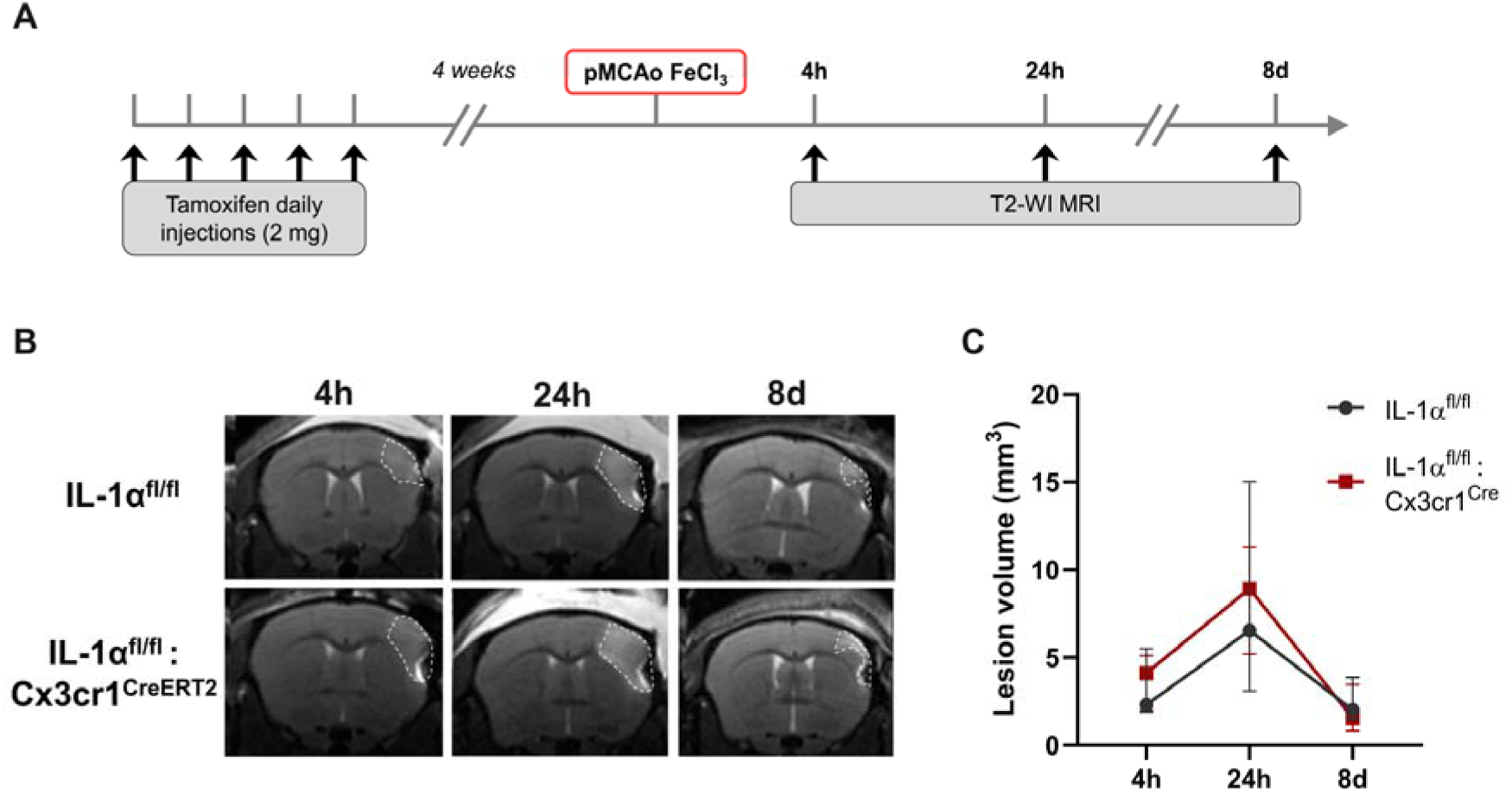
Microglial IL-1α deletion does not influence brain lesion progression up to 8 days after pMCAo. (A) Schematic representation of the experimental design. (B) Representative T2-WI at 4 h, 24 h and 8 days after pMCAo. (C) Lesion volume quantification showing no differences between IL-1α^fl/fl^ and IL-1α^fl/fl^:Cx3cr1-Cre mice (n=4/group, two-way ANOVA followed by Sidak’s post hoc test).

At 24 h after tMCAo, IL-1α^fl/fl^:Cx3cr1-Cre^ERT2^ mice showed no changes in lesion volume (46.7±15.8 mm^3^ vs. 51.5±11.9 mm^3^ in the IL-1α^fl/fl^ group, p=0.5662; Fig. 2D). The lack of effect of microglial IL-1α on brain damage was reproduced in a different tMCAo model (45 min occlusion via an incision in the external carotid artery; p>0.05 for all comparisons vs. IL-1α^fl/fl^) (Fig. S3). We also investigated whether the deletion of microglial IL-1α could affect acute immune cell responses, at 24 h after tMCAo. No differences were observed in terms of microglial density or activation in the contralateral, peri-infarct and infarct areas (p>0.05 for all comparisons vs. IL-1α^fl/fl^). Endothelial activation, measured by the expression of ICAM-1, and neutrophil infiltration were also similar in the ipsilateral brain in both genotypes (p>0.05 for all comparisons vs. IL-1α^fl/fl^); (Fig. S4).

### Effect of microglial IL-1**α** expression on long term outcome after ischemia reperfusion model

We observed that animals with microglial IL-1α deletion had a worse neurological score at 14 d (9.6±2.6 vs. 6.9±1.7 in the IL-1α^fl/fl^ control group, p=0.0228), at 21 d although not reaching statistical significance (8.1±1.7 vs. 4.9±2.0 in the IL-1α^fl/fl^ control group, p=0.1792), and up to 28 d after tMCAo (9.2±1.7 vs. 6.4±2.6 in the IL-1α^fl/fl^ control group, p=0.0229); (Fig. 4B).

**Figure 4.**
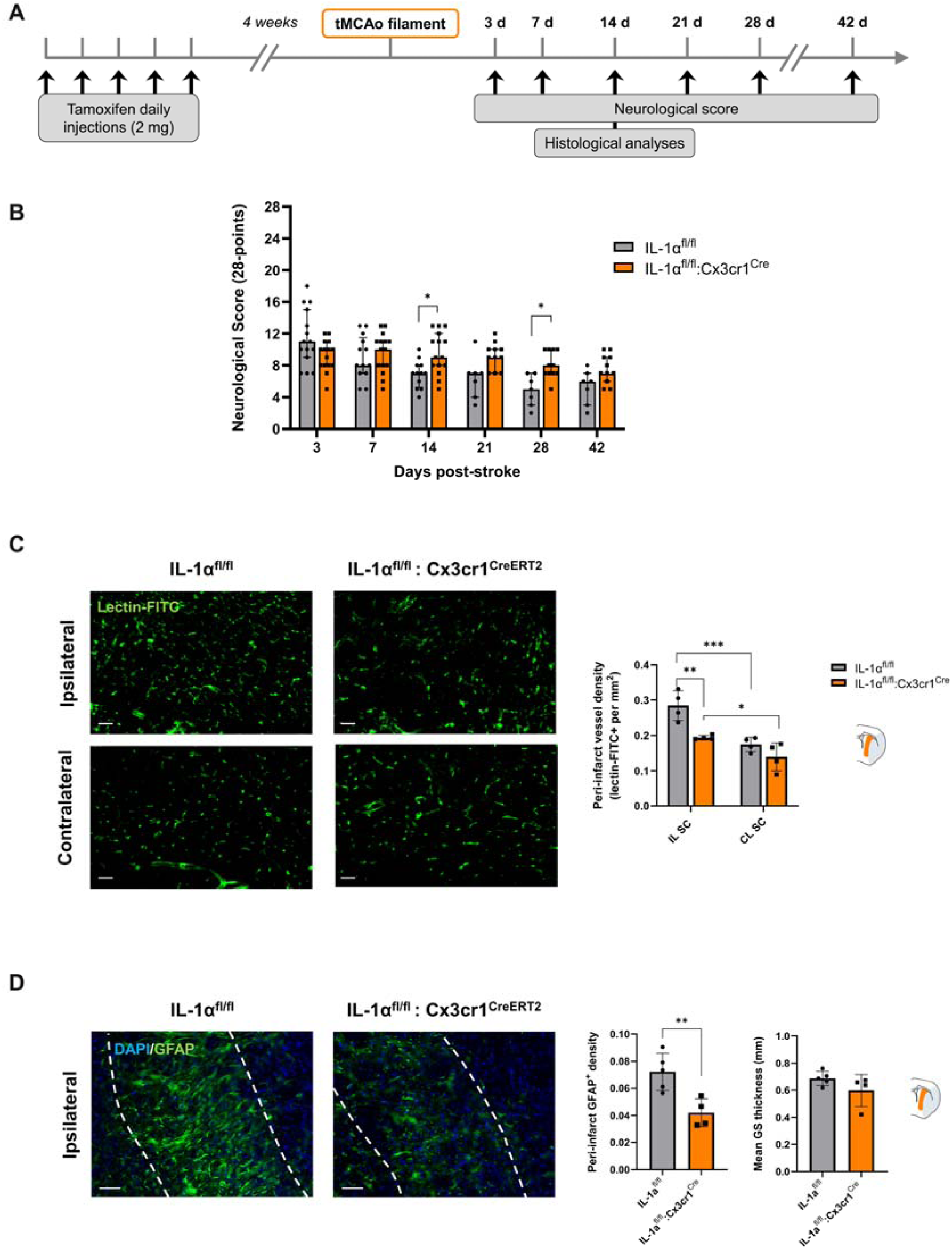
Effect of microglial IL-1α deletion on long-term stroke outcome. (A) Schematic representation of the experimental design. (B) 28-point neurological score showing significant functional deficit IL-1α^fl/fl^:Cx3cr1-Cre^ERT2^ compared to in IL-1α^fl/fl^ mice, 14 and 28 days after tMCAo (n=15-16, *p<0.05 mixed effects model (REML) followed by Sidak’s post hoc test). (C) Representative immunostaining of vascular density (lectin-FITC+) in the ipsilateral peri-infarct and corresponding contralateral subcortical brain regions in IL-1α^fl/fl^ and IL-1α^fl/fl^:Cx3cr1-Cre^ERT2^ mice, and quantification (Scale bar: 50 µm, n=4/group, two-way ANOVA followed by Sidak’s post hoc test). (D) Representative immunostaining of the glial scar (GFAP+) in the ipsilateral peri-infarct subcortical brain region in IL-1α^fl/fl^ and IL-1α^fl/fl^:Cx3cr1-Cre^ERT2^ mice, and quantification of the GFAP+ density (left) and glial scar (GS) thickness (right) (Scale bar: 50 µm, n=4-5/group, unpaired t-test, **p<0.01).

To explore the potential underlying mechanisms for this worse outcome, we performed histological analyses of the brains harvested at 14 days after tMCAo, when endogenous post-stroke neurorepair processes such as angiogenesis and neurogenesis are highly active (19).

Cortical peri-infarct vessel density was unchanged between ipsilateral and contralateral brain hemispheres in both groups (p>0.05 for all comparisons vs. IL-1α^fl/fl^; Fig. S5), whilst peri-infarct subcortical vessel density was enhanced in both IL-1α^fl/fl^ and IL-1α^fl/fl^:Cx3cr1-Cre^ERT2^ mice by 1.6-fold (p=0.0008 vs. contralateral) and by 1.4-fold (p=0.0269 vs. contralateral), respectively (Fig. 4C), which is in line with the ischemic lesion primarily affecting subcortical areas in this model (Fig. 2D).

Furthermore, we observed that IL-1α^fl/fl^:Cx3cr1-Cre^ERT2^ mice had a 4.9-fold decrease in the peri-infarct subcortical vessel density compared to that of IL-1α^fl/fl^ (p=0.0025; Fig. 4C). Along with a decreased vessel density at 14 days post-stroke, mice with microglial IL-1α depletion showed a reduced density of reactive astrocytes, measured by the GFAP+ area in the subcortical peri-infarct area (p=0.0085 vs. IL-1α^fl/fl^ mice; Fig. 4D).

Overall, microglial IL-1α deletion led to reduced vascular density and astrogliosis, which are known to contribute to post-stroke recovery and could therefore explain the long-term functional impairment amongst IL-1α^fl/fl^:Cx3cr1-Cre^ERT2^ mice, suggesting a possible role of microglial IL-1α in post-stroke neurorepair processes.

## Discussion

In this study, we show that IL-1α and IL-1β have differential spatio-temporal expression profiles in the brain after stroke, confirming earlier observations (8) and suggesting that they might exert distinct non-overlapping roles at different stages of post-stroke inflammation. Here, we show that IL-1α is expressed exclusively by microglial cells early after stroke onset, followed by a delayed IL-1β expression by a small subset of microglia and mostly by infiltrating neutrophils. Upon stroke onset, microglia are the first resident immune responders, rapidly migrating and becoming activated at the lesion site, exacerbating tissue injury by releasing pro-inflammatory cytokines and cytotoxic factors, but also contributing to tissue repair and remodeling by engulfing cellular debris and invading neutrophils, and releasing anti-inflammatory cytokines and growth factors (20, 21). With microglia being the major source of brain IL-1α expression and considering its pivotal role in post-stroke neuroinflammation, we aimed to study the specific role of microglial IL-1α in ischemic stroke using our novel microglial-specific IL-1α deficient mouse.

A previous study showed that, whilst the ubiquitous and chronic deletion of both IL-1α and IL-1β significantly reduced ischemic brain damage, the deletion of neither IL-1α nor IL-1β alone had no effects on ischemic brain injury, suggestive of potential compensatory mechanisms (22). In the present study, microglial IL-1α deletion did not influence acute brain damage and cerebral blood flow, nor infarct lesion progression as seen by MRI up to 8 days post-stroke in a model of permanent cerebral ischemia. To enhance translatability, we also assessed acute brain damage in two different intraluminal tMCAo models in different research centers, which replicated the lack of effect of microglial IL1-α deletion on infarct size, suggesting that microglial IL-1α does not contribute to acute brain damage, despite its very early expression.

The effect of microglial-derived IL-1α on microglial function in stroke is largely unknown. However, microglial proliferation is known to be the main source of microgliosis after ischemic stroke (23) and, in this regard, *IL1R1* and *IL1A* have been identified as key genes involved in microglial replenishment upon ablation in adult mice (24), while CX3CR1 has been implicated in microglial chemotaxis, microglia-mediated neurotoxicity, and microglial activation (25). With all this in mind, we investigated whether CX3CR1-Cre-mediated microglial IL-1α deletion could affect microglial functions, such as microgliosis and neutrophil engulfment, after both transient and permanent cerebral ischemia.

Microglial IL-1α ablation did not modify microglial density nor activation, neutrophil infiltration, endothelial activation, nor IL-1β expression in the brain after stroke. These results suggest that microglial IL-1α may not be involved in microglial functions nor IL-1β expression early after stroke, although future studies should include additional timepoints to elucidate potential microglial IL-1α effects on microglia in the subacute and chronic phases.

Next, we investigated the effect of microglial IL-1α expression on long term outcome after stroke in the intraluminal model, which allows us to investigate behavioral deficits beyond the acute phase. Surprisingly, despite the early expression of microglial IL-1α, which does not seem to have major effects on acute stroke outcome, its deletion was associated with a worse functional recovery from 14 days and up to 28 days after cerebral ischemia. We also observed that microglial IL-1α deficient mice had a reduced peri-infarct vessel density, suggestive of impaired post-stroke angiogenesis. Peri-infarct vascular remodeling occurs mostly during the first 2 weeks post-stroke in rodent models and is known to contribute to functional recovery (19, 26). These results could explain the behavioral deficits in microglial IL-1α deficient mice and are in line with previous findings showing that subacute IL-1α administration increases post-stroke angiogenesis in mice (10). Furthermore, impaired angiogenesis occurred along with reduced astrogliosis as seen by a decreased GFAP expression and a slightly lower glial scar thickness. Microglia comprise a highly heterogeneous and plastic cell population, displaying numerous overlapping functional states and participating in a complex crosstalk with different cell types within the neurovascular unit and infiltrating peripheral leukocytes (20). In this context, microglia, which respond earlier to injury, can modulate astrocyte activation and functions. Recently, Huang and colleagues proposed that microglia may inhibit the recruitment of neutrophils and secondary occlusions through the release of IL-1RA, which in turn modulates astrocytic CXCL1 expression (27). In line with our results, a recent study showed that activated microglia induce reactive astrocytes through the release of IL-1α, TNFα, and C1q; however, this was associated with the activation of neurotoxic reactive astrocytes (28). The role of reactive astrocytes in stroke is a subject of debate, with evidence suggesting they can both hinder and promote stroke recovery. In accordance with our results, Williamson and colleagues showed that reactive astrocytes facilitate vascular repair and remodeling, improving motor recovery after stroke (29). However, more exhaustive and integrative omics studies are warranted to gain a better understanding of the downstream effects of stroke-induced microglial IL-1α expression.

Taken together, this study identifies a critical role for microglial IL-1α in stroke, and should be therefore considered as a potential therapeutic target. Our findings suggest that microglial-derived IL-1α does not modify acute stroke outcome, despite its hyperacute expression, yet it might regulate long-term cerebrovascular responses modulating functional recovery after experimental stroke. Nonetheless, the mechanisms underlying the non-redundancy of IL-1α and IL-1β, the role of microglia and the potential link with other cell types are not fully understood. Future studies focusing on the specific role of IL-1 family members in stroke are needed to advance the understanding of stroke pathophysiology, which is essential to refine IL-1 targeted stroke interventions.

## Supplementary Information

**Additional File 1 (.docx): Figure S1.** Characterization of a conditional IL-1α mouse mutant crossed with Cx3Cr1-Cre^ERT2^ mice to induce a specific deletion of microglial IL-1α in the brain. (A) Schematic representation of the experimental design. Exon 4 of the *IL1A* gene flanked with loxP sites (IL-1α^fl/fl^), is excised upon Cre recombination induced by tamoxifen (in IL-1α^fl/fl^:Cx3cr1-Cre^ERT2^ mice), resulting in the generation of microglia-specific IL-1α KO mice. (B) Representative immunostaining of IL-1α (red), microglia (Iba1, white) and DAPI (blue) in the contralateral, ipsilateral areas at 4 h after pMCAo, showing microglial IL-1α expression abrogation upon tamoxifen administration (Scale bar: 50µm). **Figure S2.** Microglial IL-1α deletion does not influence microglial activation, neutrophil infiltration nor IL-1β expression at 24 h after permanent cerebral ischemia. (A) Schematic representation of the experimental design. (B) Representative immunostaining of neutrophils (Ly6G, green), IL-1β (red), microglia (Iba1, white) and DAPI (blue) in the contralateral, peri-infarct and infarct areas at 24 h after pMCAo (Scale bar: 50 µm). (C) Number of microglia (Iba1 positive cells), neutrophils (Ly6G positive cells), percentage of IL-1β positive microglia and IL-1β positive neutrophils in the contralateral, peri-infarct and infarct areas at 24 h after stroke. (n=6/group, two-way ANOVA followed by Sidak’s post hoc test). **Figure S3.** Microglial IL-1α deletion does not influence microglial density and activation, endothelial activation, nor neutrophil infiltration at 24 h after transient cerebral ischemia. (A) Schematic representation of the experimental design. (B) Representative immunostaining of ICAM-1 (activated endothelium, green), neutrophils (Ly6G, purple), microglia (Iba1, white) and DAPI (blue) in the contralateral, peri-infarct and infarct areas at 24 h after tMCAo (Scale bar: 50 µm). (C) Density of microglia (% of Iba1 positive cells), microglial activation (% of activated Iba1+ cells), endothelial activation (ICAM-1 positive vessels density) and neutrophils (Ly6G positive cells); n=6/group, two-way ANOVA followed by Sidak’s post hoc test, unpaired t-test.

## Declarations

### Ethics approval and consent to participate

All animal procedures were carried out in accordance with the Animals (Scientific Procedures) Act (1986), under a Home Office UK project license, approved by the local Animal Welfare Ethical Review Board, and experiments were performed in accordance with ARRIVE (Animal Research: Reporting of In Vivo Experiments) guidelines (11), with researchers blinded to experimental groups.

### Consent for publication

Not applicable

### Availability of data and materials

The raw data supporting the conclusions of this article are available from the corresponding author upon reasonable request.

### Authors’ contributions

E.L., A.G., and R.W. contributed equally to this work. E.L.: methodology, experiments execution, data analysis and interpretation, figure creation, writing and editing. A.G.: methodology, experiments execution, data analysis and interpretation, figure creation, writing and editing. R.W.: methodology, experiments execution, data analysis and interpretation, and editing. M.R.: experiments execution. B.O.: supporting information, data interpretation and editing. B.L.: experiments execution, data analysis and editing. F.W.: experiments execution, data analysis and editing. N.L.: supporting information and editing. A.D.: supporting conceptualization, supporting information and editing. D.B.: project supervision and editing. S.M.A.: project supervision and editing. G.J.B.: conceptualization, project supervision and editing. E.P.: conceptualization, project supervision and administration, and editing.

## Abbreviations

ARRIVE: Animal Research: Reporting of In Vivo Experiments
BSA: Bovine serum albumin
C1q: Complement component 1q
CBF: Cerebral blood flow
cDNA: Complementary Deoxyribonucleic Acid
CX3CR1: Chemokine (C-X3-C motif) receptor 1
CreERT2: Cre recombinase-Estrogen receptor T2 fusion protein
CXCL1: C-X-C motif chemokine ligand 1
DAPI: 4′,6-diamidino-2-phenylindole
DPBS: Dulbecco’s phosphate-buffered saline
GFAP: Glial fibrillary acidic protein
Iba1: Ionized calcium-binding adaptor molecule 1
ICAM-1: Intercellular adhesion molecule 1
IL-1: Interleukin-1
IL-1R1: Interleukin-1 receptor type 1
IL-1Ra: Interleukin-1 receptor antagonist
IL-1α: Interleukin-1 alpha
IL-1β: Interleukin-1 beta
KO: Knock out
LSCI: Laser speckle contrast imaging
Ly6G: Lymphocyte antigen 6 complex locus G6D
MCA: Middle cerebral artery
MRI: Magnetic Resonance Imaging
MSME: Multi-slice multi-echo
OCT: Optimal cutting temperature compound
PBS: Phosphate-buffered saline
pMCAo: Permanent middle cerebral artery occlusion
RNA: Ribonucleic Acid
ROI: Regions of interest
RT-qPCR: Real-Time Quantitative Polymerase Chain Reaction
T2WI: T2-weighted imaging
TE/TR: Echo time / Repetition time
tMCAo: Transient middle cerebral artery occlusion
TNFα: Tumor necrosis factor alpha
WT: Wild type

## Acknowledgements

The authors would like to thank the Bioimaging Facility, the Preclinical-MRI Facility and the Genomic Technologies Facility in the FBMH at the University of Manchester for equipment and advice in imaging.

