## Supplementary Information for "Selective deletion of interleukin-1 alpha in microglia does not modify acute outcome but regulates neurorepair processes after experimental ischemic stroke"

^‡^Current affiliation: Normandie University, UNICAEN, INSERM UMR-S U1237, Physiopathology and Imaging of Neurological Disorders, GIP Cyceron, Institute Blood and Brain @ Caen-Normandie, Caen, France.

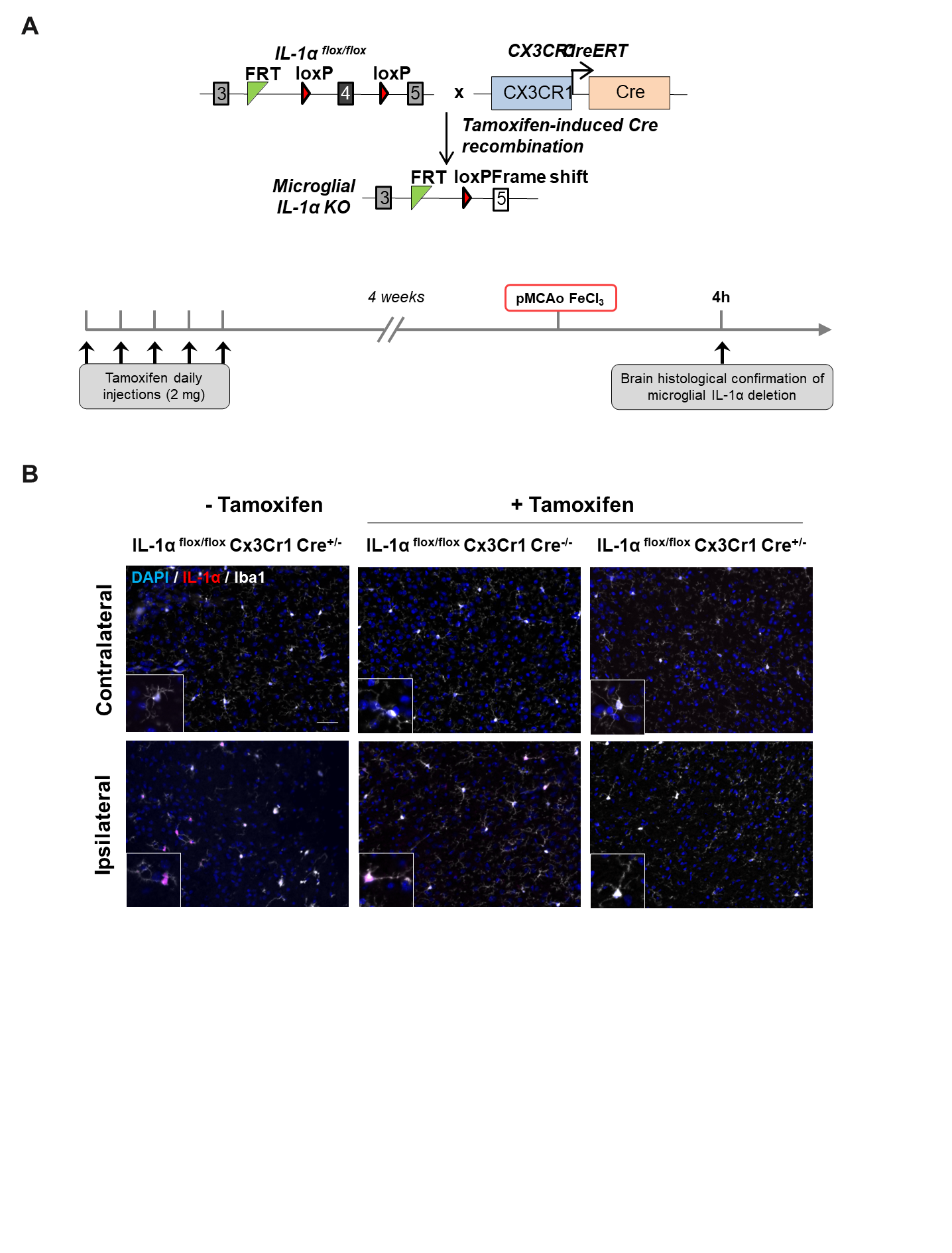

**Figure S1. Characterization of a conditional IL-1α mouse mutant crossed with CX3CR1 Cre-ERT2 mice to induce a specific deletion of microglial IL-1α in the brain.** (A) Schematic representation of the experimental design. Exon 4 of the IL-1A gene flanked with loxP sites (IL-1α^fl/fl^), is excised upon Cre recombination induced by tamoxifen (in IL-1α^fl/fl^:Cx3cr1-Cre^ERT2^ mice), resulting in the generation of microglia-specific IL-1α KO mice. (B) Representative immunostaining of IL-1α (red), microglia (Iba1, white) and DAPI (blue) in the contralateral, ipsilateral areas at 4 h after pMCAo, showing microglial IL-1α expression abrogation upon tamoxifen administration (Scale bar: 50µm).

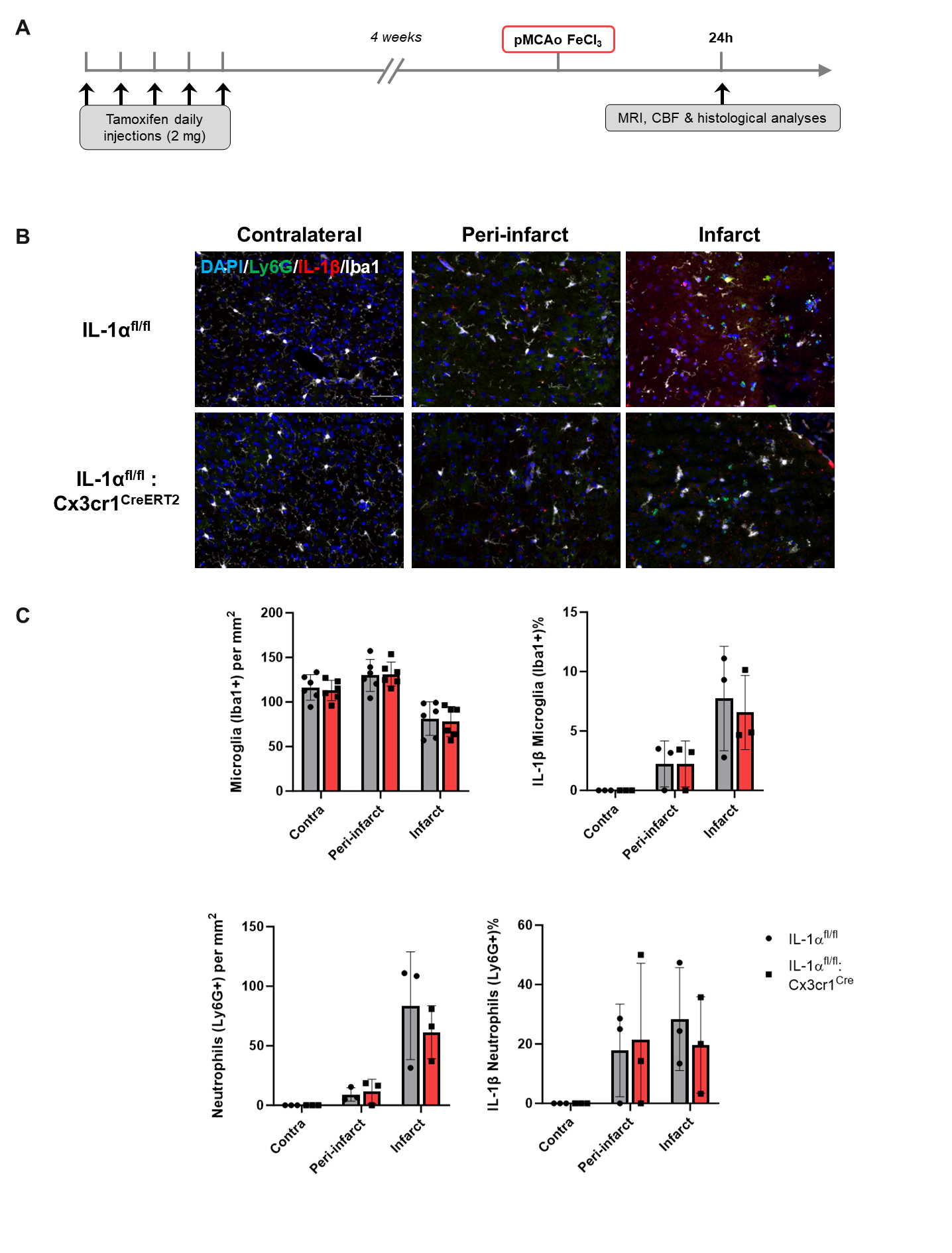

**Figure S2. Microglial IL-1α deletion does not influence microglial activation, neutrophil infiltration or IL-1β expression at 24 h after permanent cerebral ischemia.** (A) Schematic representation of the experimental design. (B) Representative immunostaining of neutrophils (Ly6G, green), IL-1β (red), microglia (Iba1, white) and DAPI (blue) in the contralateral, peri-infarct and infarct areas at 24 h after pMCAo (Scale bar: 50 µm). (C) Number of microglia (Iba1 positive cells), neutrophils (Ly6G positive cells), percentage of IL-1β positive microglia and IL-1β positive neutrophils in the contralateral, peri-infarct and infarct areas at 24 h after stroke. (n=6/group, two-way ANOVA followed by Sidak’s post hoc test).

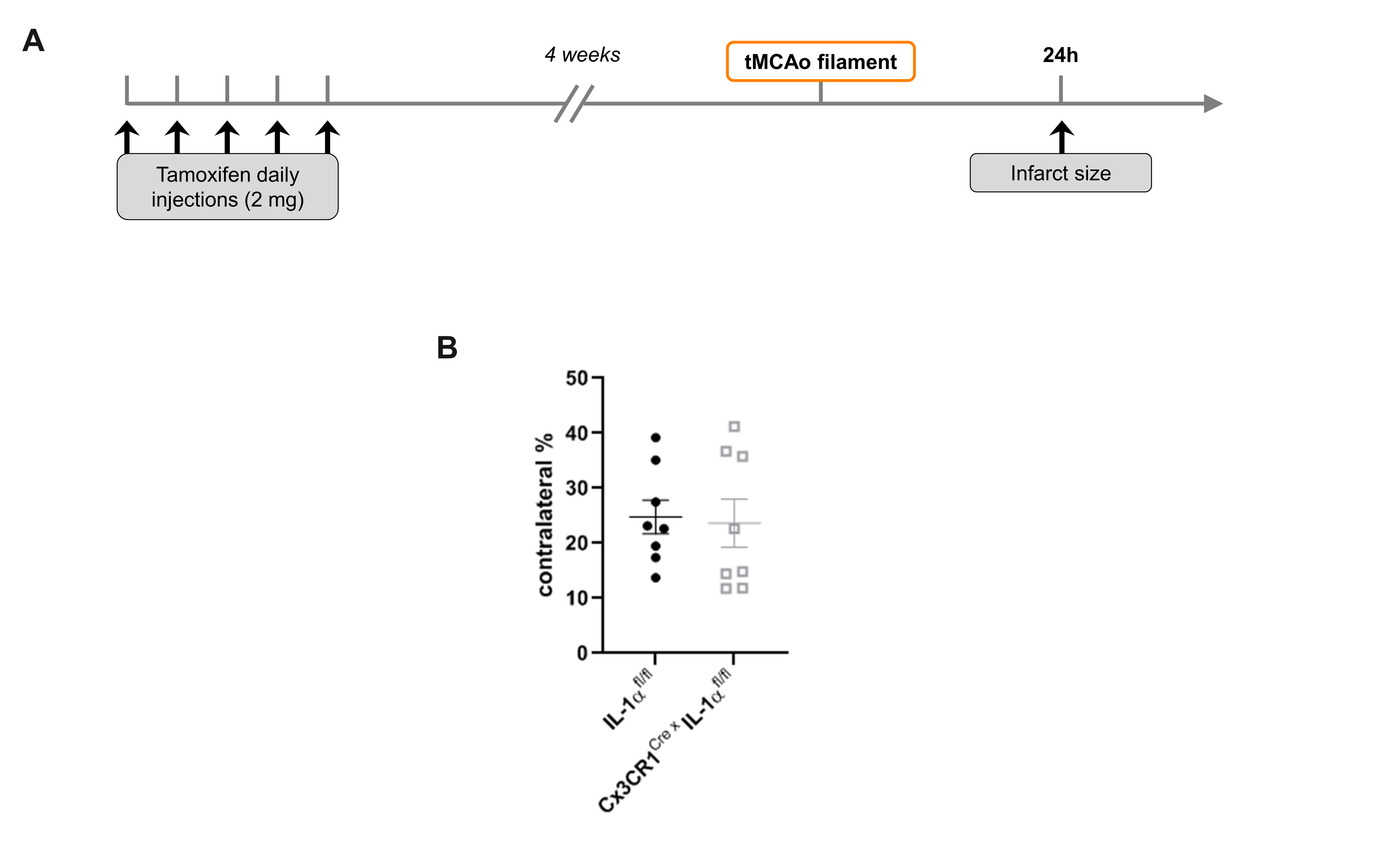

**Figure S3. Microglial IL-1α deletion does not influence brain damage at 24 hours after transient cerebral ischemia.** (A) Schematic representation of the experimental design to study acute outcome after transient stroke. (B) Infarct percentage at 24 h after tMCAo (45 min occlusion, filament through external carotid artery, ECA) in IL-1α^fl/fl^ and IL-1α^fl/fl^:Cx3cr1-Cre^ERT2^ mice (n=8/group, XX).

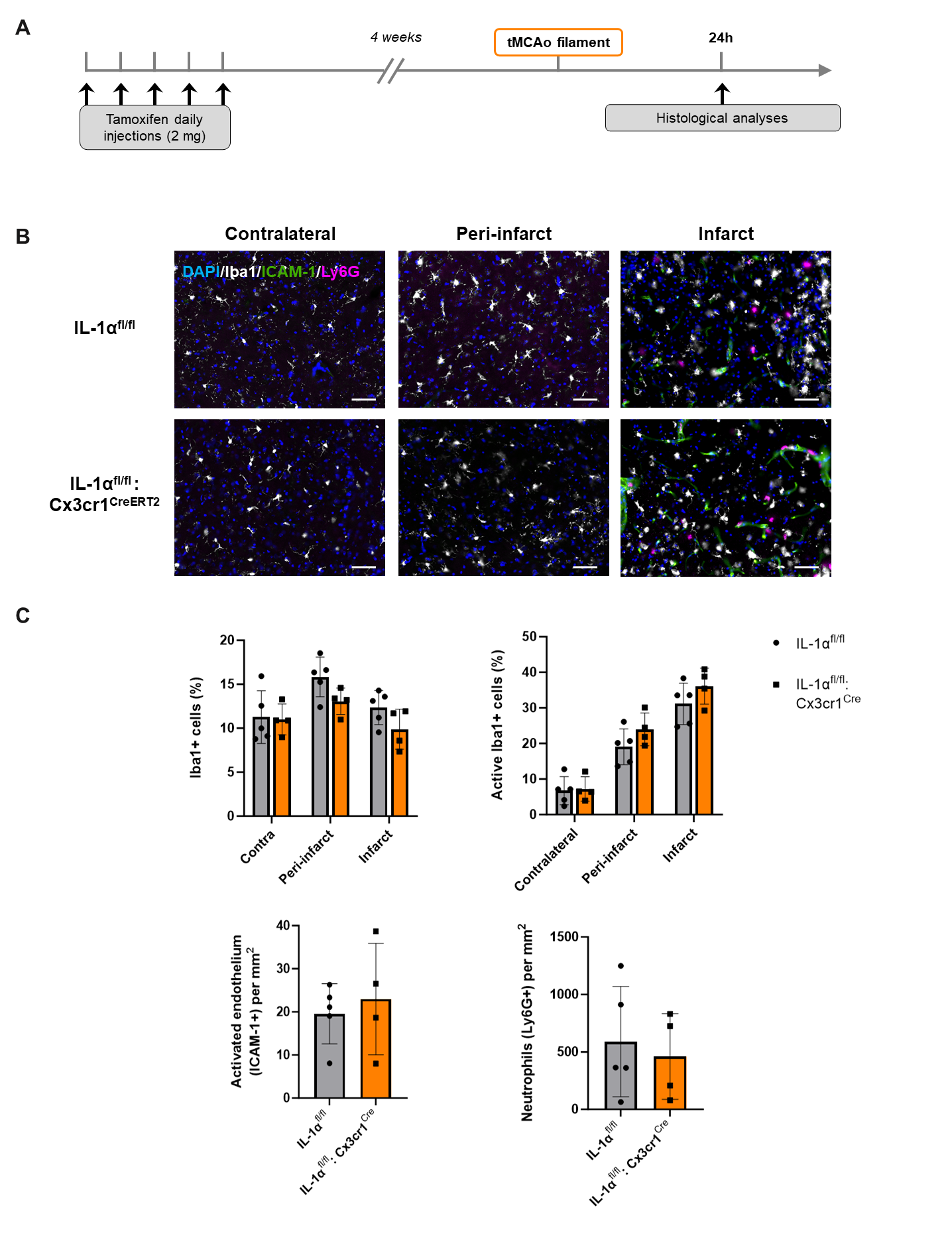

**Figure S4. Microglial IL-1α deletion does not influence microglial density and activation, endothelial activation, nor neutrophil infiltration at 24 h after transient cerebral ischemia.** (A) Schematic representation of the experimental design. (B) Representative immunostaining of ICAM-1 (activated endothelium, green), neutrophils (Ly6G, purple), microglia (Iba1, white) and DAPI (blue) in the contralateral, peri-infarct and infarct areas at 24 h after tMCAo (Scale bar: 50 µm). (C) Density of microglia (% of Iba1 positive cells), microglial activation (% of activated Iba1+ cells), endothelial activation (ICAM-1 positive vessels density) and neutrophils (Ly6G positive cells); n=6/group, two-way ANOVA followed by Sidak’s post hoc test, unpaired t-test.

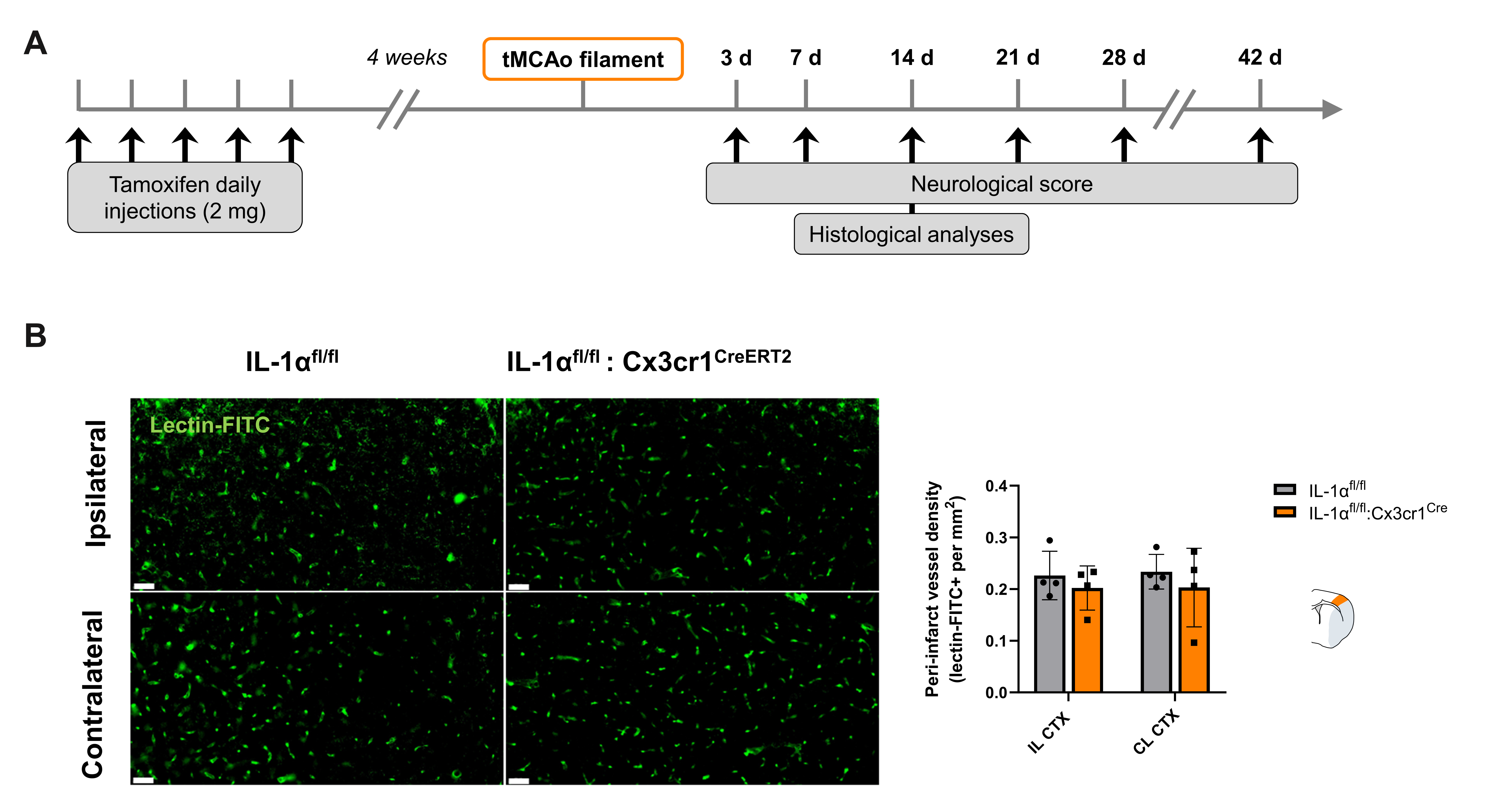

**Figure S5. Microglial IL-1α deletion does not affect cortical peri-infarct vessel density at 14 days post-stroke**. (A) Schematic representation of the experimental design. (B) Representative immunostaining of vascular density (lectin-FITC+) in the ipsilateral peri-infarct and corresponding contralateral cortical brain regions in IL-1α^fl/fl^ and IL-1α^fl/fl^:Cx3cr1^ERT2^ mice, and quantification (Scale bar: 50 µm, n=4/group, two-way ANOVA followed by Sidak’s post hoc test).
